# Real-time affect decoding in the amygdalo-hippocampal circuit from dynamic auditory signals

**DOI:** 10.1101/2025.05.09.652704

**Authors:** Florence Steiner, Sascha Frühholz

## Abstract

Affect decoding from auditory signals requires the temporal tracking and neural processing of dynamic sound patterns, such as in affective speech. Affective speech is commonly expressed to maximize its emotional impact and its neural decoding in integrated medial limbic circuits of recipients. Here we examined how affective speech, which was live produced by speakers to maximize amygdala-hippocampal connectivity in listeners, can evoke significant intralimbic and cortico-limbic affect decoding mechanisms. Aggressive and joyful affective speech that was produced based on real-time feedback of amygdala-hippocampal couplings in listeners increased intralimbic connectivity as well as activity in auditory cortical nodes as parts of a broader affective sound processing network. Neural time courses in the auditory cortex also correlated with acoustic patterns of adaptive speech indicative of a communicative speaker-listener coupling. Affective speech can thus meaningfully influence limbic circuit synchronizations with a specific significance of the amygdala-hippocampal circuit for affect decoding from auditory signals.

## INTRODUCTION

Affective signaling and communication play a significant role in human social behavior. For affective speech, a multitude of emotional cues are conveyed through prosodic voice intonations, providing valuable insights essential for navigating and adapting to social situations (1). Humans can decode emotions solely based on affective prosodic cues (2), which makes the paralinguistic speech dimension a powerful channel for affective communication. The neural core system that processes such affective voice signals expressed in speech encompasses a large area in the bilateral auditory cortex (AC) extending over low and high-order AC (3, 4), the inferior frontal gyrus (IFG) (5–7), and several areas of the limbic system (8). Important limbic brain areas for affect decoding comprise the amygdala (AMY) and closely connected areas, such as the ventral striatum and the hippocampus (HC) (8). These cortico-limbic brain systems are involved in different processes necessary for affect decoding from speech, including the extraction of acoustic features and their analysis, tracking these features over time and identifying meaningful relations among them, and evaluating these integrated acoustic patterns in relation to previous emotion-related knowledge (3, 7).

Within the brain network involved in affect decoding from speech, the limbic system represents a key component. Especially the significance of the AMY in affective speech processing is relatively well-established (6, 9), with an important role in decoding affect from dynamic speech (7). Unlike for the AMY, the involvement of the HC and especially the interplay between the AMY and HC have so far not received much attention. This is somehow surprising for several reasons. First, the AMY and HC lie in close anatomical proximity to each other (8). They are both situated in the bilateral medial temporal lobe with the AMY located immediately anteriorly to the HC, which in turn extends longitudinally further back. Second, there is dense structural interconnectivity between the AMY and HC (10–12). Third, both regions show functional similarity. Specifically, there is a functional distinction along the longitudinal hippocampal axis where the posterior HC (pHC) seems to be involved in more cognitive functions, and the anterior HC (aHC), that is directly bordering the AMY, seems instead rather engaged during emotional processes (12–14). Thus, the functional similarity with the AMY primarily applies to the aHC, which is also reflected in their anatomical interconnectivity, as only the aHC but not the pHC seems to share dense bidirectional anatomical connections with the AMY (10, 12).

Besides their close anatomical connection, the AMY and HC have also been found to functionally interact closely during a variety of processes. For example, the AMY has an enhancing influence on episodic memory consolidation in the HC (15). Furthermore, inputs from the AMY to the aHC also seem to have a modulating influence on social approach behavior (16, 17). Reversely, the HC supplies contextual information to the processing of emotional stimuli in the AMY for recollecting past experiences (18) or stored templates of affective vocalizations (8). Additionally, the HC also seems to specifically be engaged in processing social emotions like admiration and compassion (19). Apart from being involved in processing emotional and social situations, the HC also plays a role in processing and tracking the temporal aspect of sensory information in general (20). While temporal processing is important for all sensory modalities, it has a special relevance for auditory information as it usually develops over time and the HC supports temporal information tracking (21).

Apart from these AMY-HC interactions for a variety of cognitive processes, the HC has also received more attention regarding its involvement specifically in auditory processing in recent years (21, 22). The HC is affected by noise on several physiological levels, including neurotransmission, neurogenesis, and electrophysiology (22). Additionally, the HC and its communication with the AC seem to be altered in auditory disorders (tinnitus, hearing loss) (22). These interactions are also undermined by direct projections from the hippocampal CA1 subfield to the auditory cortex, which likely reflect the direct information exchange between those regions (23). There are also many indirect connections between the auditory system and the HC and especially subcortical routes allow for a fast relay of early auditory information (21, 24).

This altogether seems to point to the functional significance of HC activity and HC coactivations with the AMY for some important affect decoding functions, especially for auditory emotions. Accordingly, HC activity is frequently reported in studies on auditory emotions encoded in voice signals (8), especially for socially more complex or ambiguous affective tones like guilty, proud, bored (25), ironic, doubtful (26), or surprised (27). Other studies found HC activity during discrimination of negative (28) and positive affective voices (29) as well as during human affective speech compared to synthetic speech processing (26) or for whispered affective speech (30). These observations point to the relevance of contextual factors during affective speech decoding in the AMY-HC circuit, which is often also observed in auditory contextual conditioning (31).

Summarizing previous evidence, the HC seems to integrate dynamic temporal auditory information and relatedly should support the AMY in the real-time evaluation of affective speech in socially and dynamically more complex contexts. Especially in real-life contexts, affective communication is based on dynamic speaker-listener interactions with mutual communicative adaptions. Neural processes involved in successfully participating in such a dynamic interaction include evaluation of emotional, social, and temporal contexts as well as temporal integration of communicative auditory and affective information. However, many previous studies on affective speech (3) used non-dynamic and non-adaptive experimental designs, with vocal emotions usually presented as pre-recorded stimuli with short durations. These studies might therefore have missed some of the neural effects that affective speech would have in more real-life communicative situations, and this might explain inconsistent findings about the involvement of the HC in affective speech decoding until now (8). Introducing more ecologically valid experimental designs of affective communication might better determine the importance and significance of the AMY-HC axis in affective speech decoding.

We accordingly developed an innovative experimental design for real-time neuroimaging that connects speakers and listeners in a closed-loop setup where speakers can dynamically modulate their affective speech to actively influence listeners’ neural affect decoding mechanisms in real time involving significant temporal dynamics (Fig. 1). We specifically focused on the functional connectivity in the AMY-aHC axis to further elucidate and potentially solidify and specifically determine the role of the HC in affective speech processing. An important issue in neurofeedback studies, which is related to the specificity of the investigated neural mechanisms, concerns the inclusion of appropriate control or baseline conditions (32). The control condition is relevant to determine the specificity of the target neural mechanisms controlled by the feedback setup and is used as a comparison condition. Typical options for control conditions are using neurofeedback signals from a mechanistically irrelevant brain area or using sham/random neurofeedback signals. We tested both possibilities in the piloting phase of the experiment, but these conditions turned out to be very confusing for the speakers as they considerably impaired the continuous and fluent production of affective.

**Figure 1.**
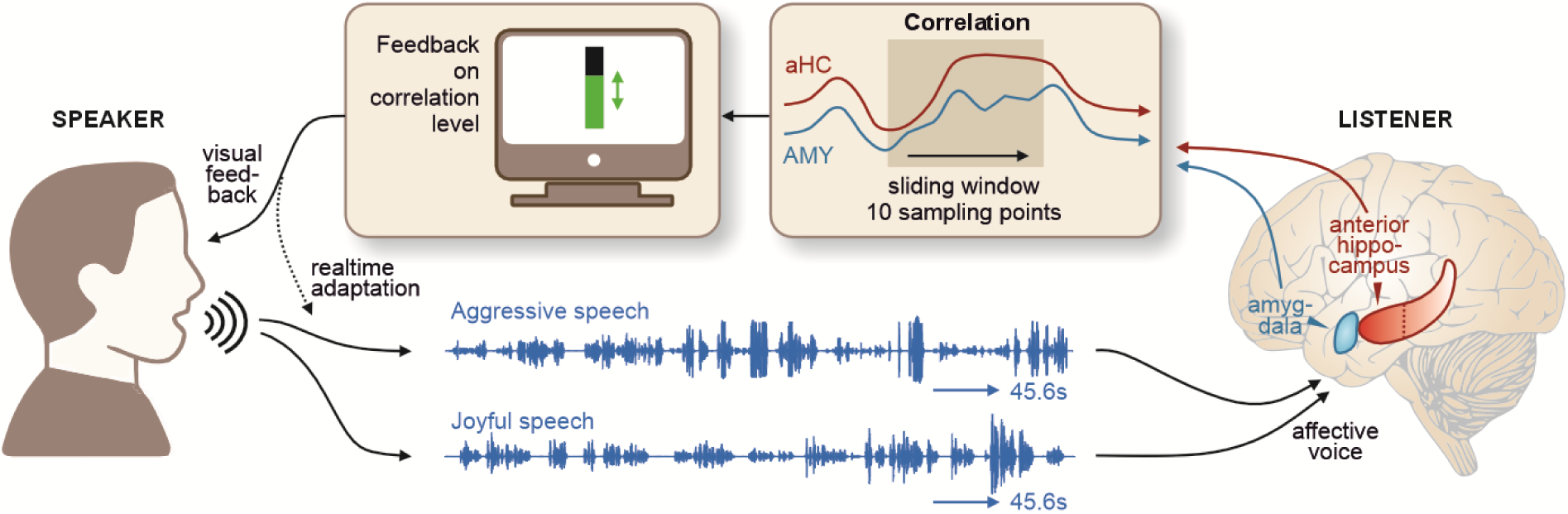
Closed-loop setup linking a speaker and listener in adaptive affective speech communication. Speakers continuously modulated their affective speech (aggressive or joyful) in response to the level of connectivity between the amygdala (AMY) and the anterior hippocampus (aHC) quantified in listeners’ brains while they listened to the speakers’ voices. The AMY-aHC connectivity signal was quantified in real time by using a sliding window across the 10 last sampling points and visually displayed to speakers as a continuously updated bar graph representing the level of correlation at each scanning time-point. The speakers’ task was to modulate their affective speech intonations with the aim to introduce real-time adaptions in order to maximize the AMY-aHC connectivity in listeners.

We therefore designed our experiment including two main experimental conditions and one control condition. First, we compared neural connectivity in the AMY-aHC axis during this real-time dynamic speech, closely resembling real-life affective speech dynamics, with neural connectivity while listening to affective speech that speakers produced without receiving feedback from listeners’ brains as the control condition. The latter condition was identical to the adaptive dynamic speech condition (same speakers, some affective tones) but without the possibility of individually adapting the affective speech to a listener’s neural reactivity. Second, we included two main experimental conditions, such that speakers adapted their affective speech either for feedback signals originating from the left or from the right AMY-aHC connectivity in separate experimental blocks. Including these two main experimental conditions allowed us to determine the specificity of neurofeedback effects in the left or the right medial infralimbic brain system. We hypothesized that the AMY-aHC connectivity can be specifically targeted and maximized with dynamic and adaptive affective speech. We also expected that increased AMY-HC connectivity would elicit increased activity in the broader brain cortical and subcortical network for affective speech decoding (3).

## MATERIALS AND METHODS

The general experimental setup included speakers vocalizing affective intonations superimposed on pseudo-speech sentences as well as listeners of these affective speech utterances. Listeners’ brain activity and limbic connectivity was quantified in real time by functional magnetic resonance imaging (fMRI) and served as a feedback signal to the speakers. Specifically, speakers were presented with listeners’ real-time connectivity between the amygdala (AMY) and the anterior hippocampus (aHC), which represents a specific component in the neural affective speech processing system. By modulating their speech intonations in adaptation to this neural target signal, speakers were asked to maximize processing in the affective speech processing system of listeners.

### Participants as listeners for affective voice perception

A total of 25 healthy volunteers (10 female, mean age 28.4y, SD 6.4) participated in the experiment as “listeners”. They were asked to listen to samples of affective speech that “speakers” vocalized in real time and in direct adaptation to the listeners’ neural responses. Four additional participants had to be excluded from analysis for various reasons, like technical or procedural errors during the experiment resulting in missing acoustic or neural data or excessive movement. The listening participants had normal hearing abilities and normal or corrected-to-normal vision. No participant presented a neurological or psychiatric history. All participants gave informed and written consent for their participation in accordance with the ethical and data security guidelines of the University of Zurich. The experiments were approved by the Swiss governmental ethics committee.

### Vocal speakers for affective voice production

All 10 vocal speakers involved in the experiment were advanced acting students from the Zurich University of the Arts and trained actors (5 females; mean age 27.6y, SD 4.1). Through their acting studies, these actors were experienced and trained in expressing and portraying affective states, and they had advanced vocal training. Thus, they represented the ideal speakers for our experimental setup as it was necessary that the speakers were in full control of their vocal range, able to introduce a variety of vocal changes, and could believably and authentically transport the respective emotions with just their voices. Additionally, they were adept at learning text by heart, which was important to cite the text fluently and fully pay attention to only the listeners’ AMY-aHC connectivity.

### Experimental conditions

The pseudo-sentences were spoken in three different experimental conditions. In two conditions the speakers received real-time dynamic feedback presented as a vertically changing bar graph and based on a target signal extracted from listeners’ brains to which they adapted their voices. The target for the feedback extraction was the connectivity between the ipsilateral AMY and aHC. One condition focused on the connectivity feedback from the left hemisphere (OnLfeed condition) and was the main experimental condition and the second condition focused on the connectivity feedback from the right hemisphere (OnRfeed condition) and served as a control region condition. Both conditions together are referred to as the Onfeed conditions and the vocalizations are referred to as “feedback-based adaptive speech”.

The third condition served as the baseline condition. Speakers received no live feedback but were instead presented with a static display of the feedback bar at medium height carrying no dynamic information about the listeners’ brain connectivity (Offfeed condition). In this condition, speakers were thus not able to adapt their voice individually to the listeners’ neural reactions and vocalizations will accordingly be referred to as “non-adaptive speech”.

### Experimental design

Each speaker vocalized each of the three experimental conditions once in a joyful and once in an aggressive vocal tone. Thus, the experiment had a 3×2 factorial design, including the factors “speaking mode” (three levels: OnLfeed, OnRfeed, Offfeed) and “emotional tone” (two levels: aggressive, joyful). Each combination of condition and emotional tone was presented in a separate block during the fMRI experiment resulting in six blocks per speaker. All speech conditions for one listening participant were produced by two speakers (one male, one female) randomly selected from the total ten speakers. While ensuring an equal distribution across all participants, we randomly assigned the speakers to the listeners. Thus, each listener was presented with a total of 12 vocalization blocks (3 speaking modes 2 emotional tones 2 speakers). Each of the twelve blocks included an alternating sequence of six resting periods with no auditory stimulation (each 20.4s = 17TRs) and five speech periods (each 45.6s = 38TRs). The order of the twelve blocks was random and counterbalanced across participants. In all twelve blocks, participants listened to live speech of the actors that was either adapted to the neural target feedback or not.

We introduced the factor “speaking mode” because we are interested in the influence of dynamic and individually adapted affective communication in comparison to affective speech that is not adapted to a listener’s reaction as it was done in many previous studies in the research field (1, 3). The individually adapted speech is an ecologically more valid design feature in relation to many real-life and “live” settings with affective communication being an adaptive and interactive process. Comparatively, the approach frequently used in previous affective speech studies included using recorded affective speech during the experiment (3).

In general, in real-time fMRI neurofeedback studies where brain activity is regulated by the participants inside the scanner themselves, an appropriate baseline condition needs to be chosen as a comparison to the blocks where the brain signal is actively modulated. Commonly, studies compare brain regulation against non-regulation blocks. Therefore, we chose two conditions that differed in the possibility for the speakers to dynamically modulate their speech intonations and to regulate the listening participant’s neural target signal. The non-adaptive condition (Offfeed) thus served as the baseline condition where no regulation of and adaption to the listener’s brain signal was possible. Apart from the non-adaptive nature, however, the non-adaptive condition was exactly the same as the feedback-based adaptive conditions because it included the same speakers on the same day, the same expressed emotions, microphone set-up, and modulation of vocal features; the latter, however, was introduced in response to an imagined rather than a real social interactor.

Additionally, we wanted to introduce another control feedback condition. A less commonly used baseline condition is using an experimentally unrelated control region/target for quantifying a random neurofeedback signal. While we assessed this possibility in a piloting experiment, the randomness of the neurofeedback signal resulted in confusion for the speakers regarding the task to modulate their voice in real time. During voice modulations adapted to the neurofeedback of the neural target speakers learned appropriate strategies to maximize the neural target signal in each individual listener. However, those strategies were not applicable for the unrelated control regions and the random feedback thus led to confusion and the unlearning of the acquired strategies. As a compromise, we thus decided to use our planned target (AMY-aHC connectivity) in both hemispheres separately. Thus, AMY-aHC connectivity from each hemisphere was extracted and used individually so that one feedback-based adaptive condition focused on the connectivity in the left hemisphere (OnLfeed) and one condition in the right hemisphere (OnRfeed). This allowed us to compare the influence of affective speech intonations adapted to the main experimental feedback target with speech adapted to a different “control” target that did not confuse our speakers. Additionally, this approach provided the chance to explore potential lateralization effects.

### Task of the fMRI participants (affective speech perception)

Listeners were instructed that they will hear affective voices speaking in a pseudo language without semantic meaning, and that they should carefully and attentively listen to the affective quality of the voice. The listening participants were unaware of the different conditions, especially that the setup included live speakers and that their AMY-aHC connectivity was being modulated during the feedback-based adaptive conditions by modulations of the speakers’ affective expressions. To assess whether participants attentively listened to each speech period, we asked questions after each block (“Which emotion did you hear?”, “Was it a male or female voice?”, “How attentive were you from 1 to 5?” (1 being the lowest)) that ensured that they were focused on listening to the speech intonations.

### Task of the speakers (affective speech production)

The speakers were asked to upregulate the activity of the AMY-aHC connectivity of listeners by dynamically modulating their affective speech intonations according to several different features, such that they introduced vocal variations that potentially drive a signal increase in listeners’ AMY-aHC connectivity. The neural target signal was represented by visual analog feedback in the form of a vertical bar graph, and the task was to increase the bar in the positive direction as much as possible.

The actors were extensively trained before the experiment on emotional intonations and how specific emotions prototypically sound like. The actors also learned the pseudo-speech sentences before the experiment to enable free and fluent speech performance in the live condition. With the training we ensured that actors would not simply introduce significant differences in speech rate and voice intensity (loudness), but rather to use their full vocal potential and introduce vocal modulations along several dimensions of the voice acoustic space (timbre, pitch, harmonics-to-noise ratio as well as vocal variations on these features). The actors were also asked to imagine specific situations for each emotional tone (joyful: reminisce about a happy memory, proclaim the adoration for something, etc.; aggressive: scold someone, be irritated about an annoying person) during vocalizations. Using this repertoire of vocal and mental strategies they were asked to elicit an increase in the AMY-aHC connectivity along the affective tone expressed. Since each listeners’ AMY-aHC connectivity reacted slightly different, actors were asked to learn the best strategy for each listener within the constraints of the target emotions and vocal strategies. The ability of actors to modulate their voice along the target emotions was quantified by the post-experimental ratings and the acoustic voice analyses.

For the non-adaptive condition, actors were instructed to speak to an imaginary other person and to modulate their speech to imagined responses of the person, which is also a common part of vocal acting training in general. Although these speech intonations would be acted and not deeply “felt” by these actors, we have recently shown that these “acted” vocal emotions can be classified as being like the target emotion and include bodily responses in speakers that resemble real emotions (1–3).

### Experimental affective voice stimuli

As experimental stimuli for the affective speech, we used two pseudo-sentences consisting of pseudo-words that followed general linguistic and phonetic rules, thus resembling normal speech but without carrying any semantic meaning. The sentences were similar to previously used pseudo-sentences (1, 3, 4). To prevent speaking pauses and to allow flexibility in the modulating performance of speakers, the sentences missed punctuation. As we purposely chose meaningless words, the focus of the experiment was entirely on the acoustic properties of the speech instead of the content. Speakers were instructed to speak these pseudo-sentences continuously without pauses for a duration of 45.6s in a neutral and two different affective tones (i.e. aggressive and joyful). They vocalized the affective vocal tones live during the main fMRI experiment and the neutral tone was recorded with the same microphone setup after the main experiment for use in a post-experimental perceptual rating. For the perceptual rating and further acoustic analyses, all live speech periods produced during the main experiment were simultaneously recorded and later cut to exactly 45.6s (the duration of the speech periods). The recordings were 16-bit recordings with a sampling rate of 44.1kHz.

### Experimental setup and procedure

The experimental setup included several major parts. The imaging session started with the recording of an anatomical and a few functional images of the listening participant’s brain. We then performed a functional voice localizer sequence with the listeners. During the voice localizer, the anatomical and functional images were analyzed using the BrainVoyager software (v2.8; RRID:SCR_013057; brainvoyager.com/products/brainvoyagerqx.html) and the following steps were applied to the anatomical image: intensity inhomogeneity correction, brain extraction, and transformation to MNI space. The transformed anatomical image was then co-registered with MNI-masks of the bilateral AMY and aHC used for subsequent feedback extraction. Next, the native-to-MNI-transformation parameters were used for backwards-transformation of the AMY and aHC masks to native space. Finally, the anatomical and functional images were co-registered to ensure correct feedback extraction during the real-time fMRI experiment.

For the main experiment, the speakers performed their affective vocalizations live into a microphone that was directly linked to the headphone of listeners inside the fMRI machine. The microphone was positioned in an anechoic chamber setup located in a room near the MR scanner and connected to the experimental computer via cable connections where all live speech periods were recorded (see Fig. 1). The main experiment comprised 12 experimental real-time blocks, four blocks per speaking condition. Two speaking conditions included dynamic real-time feedback from the neural target (intrahemispheric AMY-aHC connectivity, one condition in either hemisphere) to which the speakers adapted their speech intonations accordingly (feedback-based adaptive speech or Onfeed conditions). Thus, during eight of the twelve blocks we continuously sampled brain activity from the whole brain and the neural targets. The sampled target signal was then fed back to the speakers in real time via a dynamic vertical visual scale. In the remaining four blocks (non-adaptive speech or Offfeed condition), the speakers were presented with the same vertical visual scale but instead of dynamic real-time feedback, the visual scale was set to a fixed height at the middle of the scale and thus providing no listener neural feedback information.

After the fMRI experiment, we asked the listening participants to perform a post-experimental rating. They rated each of the speech intonations they had heard during the fMRI experiment on the perceptual dimensions of emotional arousal and valence (see below).

### Functional voice localizer scan

To localize voice-sensitive cortical regions, we used a standard functional voice localizer scan procedure (5, 6). The stimulus material consisted of forty 8s sound blocks, including 20 human speech and non-speech vocalizations (vocal sounds) and 20 non-human vocalizations and sounds (non-vocal sounds: animal vocalizations, artificial sounds, natural sounds). Sound blocks were presented at approximately 70 dB SPL and in random order separated by 4s silent breaks.

### Brain image acquisition

We recorded functional imaging data on a 3 T Philips Achieva Scanner equipped with a 32-channel receiver head-coil array and using a T2*-weighted gradient echo-planar imaging (EPI) sequence including parallel imaging (SENSE-factor 2) with the following parameters: 23 sequential axial slices, whole brain, TR/TE: 1.2s/30ms, FA 82°, slice thickness 3.0mm, gap 1.1mm, FOV 240×240 mm, acquisition matrix 80×80voxel, voxel size 3mm3, axial orientation. Structural images were acquired using a high-resolution magnetization prepared rapid acquisition gradient echo (MPRAGE) T1-weighted sequence and had 1-mm isotropic resolution (TR/TE: 6.73/3.1 ms, voxel size 1 mm3, 145 slices, axial orientation). Acoustic speech stimuli were presented via MR-compatible headphones (MR Confon®, Magdeburg, Germany) with a noise-reduction system at an approximate 70 dB SPL. Acoustic loudness during the live conditions was adjusted for each participant and speaker prior to each experimental acquisition and monitored during the acquisition to keep the volume at an approximately constant level.

### Preparation of amygdala and anterior hippocampus masks

For both regions, we chose ROI masks from the Harvard-Oxford subcortical structural atlas (RRID:SCR_001476; https://identifiers.org/neurovault.image:1700), a probabilistic atlas in MNI-space based on the segmented brains of 37 humans. In probabilistic atlases, it is possible to choose a probability threshold of each voxel belonging to a certain ROI, which is based on the overlap of ROIs across all subjects used to create the atlas (i.e. the higher the threshold, the more likely it is that a voxel is part of a ROI). From the three available probability thresholds, we chose a threshold of 25% which represents a compromise between no threshold and a rather rigid threshold of 50%. This means voxels towards the boarders of each ROI are included but there is still some degree of control to not include surrounding non-ROI voxels. To ensure a proper separation of values belonging either to voxels of the AMY or the HC, we edited the atlas ROI masks by deleting the bordering voxels at the intersection of the AMY and HC. Additionally, because we were only interested in signal from the anterior part of the HC, we deleted the posterior part oriented approximately at the uncal apex landmark and its approximate coordinate in MNI-space of y=-21 mm (7). As both steps were done manually, this resulted in slightly differing numbers of voxels between the hemispheres. Thus, our masks included 314 voxels for the AMY and 512 voxels for the aHC in the left hemisphere and 361 and 564 voxels for the right AMY and aHC, respectively (Fig. 2A).

**Figure 2.**
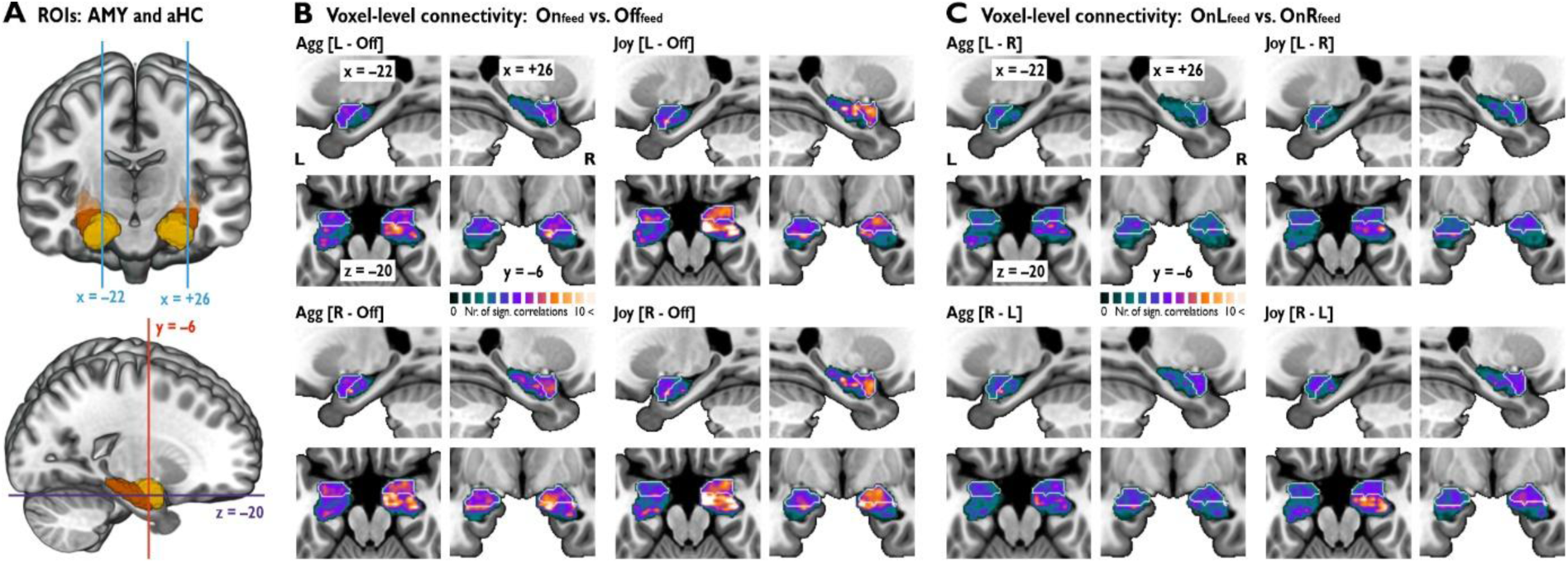
Amygdala-hippocampal connectivity is increased by adaptive affective speech. (A) Bilateral amygdala (AMY; yellow) and anterior hippocampus (aHC, orange) anatomical ROIs. The slice coordinates refer to the slices shown in panel B and C. (B) Correlation analysis between the voxels of the ipsilateral AMY and aHC. Correlation maps separately shown for both affective tones (aggressive/agg in left panels, joyful/joy in right panels) for the difference between Onfeed and Offfeed speech (comparison for OnLfeed in upper panels and for OnRfeed in lower panels). (C) Correlation maps for the difference between OnLfeed and OnRfeed speech (comparison L vs R in upper panels and R vs L in lower panels). Correlation analysis threshold p<0.05, permutation-based statistics. Higher intensity values show higher number of significant connections from one voxel to all voxels of the other ROI. Slice coordinates of each plane are the same for all contrasts and their location is shown in A. On all slices, the AMY is delimited by a white line. See also Supplementary Fig. S1.

### Online functional brain data analysis and feedback extraction

In the adaptive speech blocks during the main fMRI experiment, the AMY-aHC connectivity of the listeners was continuously quantified and fed back to the speakers. We used the TurboBrainVoyager software (v20.4; RRID:SCR_014175) to perform real-time fMRI data analysis to continuously extract activity from the AMY and aHC. The neural activities in the AMY and aHC were then correlated using the Matlab software (2018b; RRID:SCR_001622) as a measure of real-time functional connectivity.

Real-time functional brain data analysis included 3D head motion correction (all volumes are aligned to the first recorded volume of the first run), and temporal filtering (drift removal, linear trend confound predictor added to design matrix). For the online functional brain data analysis, we extracted AMY and aHC activity at each acquisition timepoint as the mean signal of all voxels within the ROI mask. The AMY-aHC connectivity feedback was then calculated at each timepoint as a Pearson correlation coefficient between the signal time courses of the current and the nine immediately preceding functional images of each ROI (correlation over the last 12s/10*TR). The number of timepoints included in the correlation was based on a trade-off and was determined in a piloting experiment. On the one hand, it was important to have an interval as short as possible, since the speakers were directly and continuously adapting their speech to the resulting correlation and the longer the interval, the less information the correlation inherited about the neural state at a specific timepoint. On the other hand, correlations needed a certain number of values to be reliably calculated. Thus, the correlation interval was defined as the lowest number of timepoints, with which the correlation could still be reliably calculated and was set to 10 TRs. At the beginning of each speech period, for TR 1-9 the correlation was calculated over all existing timepoints (e.g. for TR=5 the correlation was over the last 6s/5*TR). As explained above, correlations with less than ten timepoints were not considered reliable. Therefore, data acquired at the beginning of each speech period (TR<10) was considered noisy and in subsequent analyses we thus excluded these timepoints from the respective datasets used for analysis.

Before presenting the correlation values as feedback to the speakers, they were normalized as signal change in relation to a reference period of the current brain state during rest. The reference signal was calculated separately for each 45.6s speech period as the correlation of the last eleven scans in the immediately preceding resting period prior to each speech period. The reference signal was displayed as the zero point at the bottom of the visual scale at the start of each speech period. As we were only interested in positive AMY-aHC correlations, the visual feedback display was represented by a vertical bar that increased from the respective zero point in case of a positive correlation and was displayed as zero in case of a negative correlation. The range of the feedback bar was set to correlation change values of [0 0.8]. This range was based on a piloting experiment in which we determined the optimal height for visualization purposes based on the approximate range of the feedback value across the whole piloting experiment.

### Offline functional brain data analysis

Preprocessing and statistical analyses of functional images were performed with the Statistical Parametric Mapping software (SPM12, v7487; RRID:SCR_007037) implemented in Matlab. Functional data were first manually realigned to the anterior commissure-posterior commissure (AC-PC) axis, with the origin set to the AC. Then we used the spatial-adaptive non-local means (SANLM) denoising implemented in the CAT12 toolbox (12.5; RRID:SCR_019184) to achieve a more robust tissue segmentation result in the structural images, followed by motion correction of the functional images. Each participant’s structural image was co-registered to the mean functional image and then segmented with CAT12 to allow estimation of normalization parameters. Using the resulting parameters, we spatially normalized the anatomical and functional images to the MNI stereotactic space. The functional images for the main experiment were resampled into 2mm3 voxels. All functional images were spatially smoothed with an 8-mm full-width at half-maximum isotropic Gaussian kernel.

For first-level data analysis from the functional voice localizer scan, we used a general linear model (GLM), and vocal and non-vocal trials were separately modelled with a boxcar function aligned to the onset of each stimulus and a duration of 8s, which was then convolved with a standard hemodynamic response function (HRF). We accounted for serial correlations in the fMRI time series by an autocorrelation function of the order one. Motion parameters, as estimated during the realignment pre-processing step, were entered into the design as regressors of no interest. Contrast images for vocal and non-vocal trials were then taken to separate random-effects group-level flexible factorial analyses in order to determine voice-sensitive regions in both hemispheres of the AC by comparing vocal against non-vocal trials.

The first-level and second-level analyses for the main experiment were identical to those of the functional voice localizer scan, except that we modelled the brain data in the first-level analysis by using the finite impulse response (FIR) model. Using this FIR model, each of the 45.6s periods were modelled with a model order of 38 (corresponding to 38*TR = 45.6s) and a window length of 45.6s. This FIR modelling allowed us to better account for signal variation in the BOLD signal during the 45.6s speech period, which was highly expected during the live speech conditions. Contrast images were created for each of the six conditions by estimating the signal across all 38 time bins for each condition. The MNI coordinates of all significant peak voxels are listed in Supplementary Table S1.

All contrast images for the main experiment and the functional voice localizer scan were thresholded at a combined voxel-level threshold of p<0.005 and a cluster extent threshold of k>46 for the functional voice localizer and k>64 for the main experiment; The combined voxel- and cluster-level threshold resulted in a threshold of p<0.05 corrected at the cluster level. This cluster-corrected threshold was determined by using the 3DClustSim algorithm implemented in the AFNI software (AFNI_18.3.01; RRID:SCR_005927; www.afni.nimh.nih.gov/afni) including the new (spatial) autocorrelation function extension based on the estimated smoothness of functional images.

### Voxel-to-voxel correlation mapping to estimate AMY-aHC functional connectivity

To investigate the connectivity between the AMY and aHC on a more detailed scale, we performed a voxel-to-voxel correlation mapping between the two ROIs. The regions were defined in MNI space through the same masks used to extract the live connectivity feedback during the main fMRI experiment, including 314 voxels for the left AMY and 512 voxels for the left aHC and 361 and 564 voxels for the right AMY and aHC, respectively. We extracted the time courses of each single voxel within the ROI masks for each speech period in the interval TR 8-44. This interval included a post-trial period of 7.2 s/6*TR after the end of the speech period to account for any delays in the BOLD response and its relaxation to baseline. Despite that the connectivity feedback for the speakers was considered less stable under 10*TR, TR = 8 was chosen as a tradeoff between extracting an optimal amount of data for the correlation analysis while excluding early noisy activity and thereby increasing the signal-to-noise ratio. To localize the resulting connectivity patterns within the subfields and parts of the aHC and the nuclei of the AMY, we used probabilistic cytoarchitectonic maps as implemented in the SPM Anatomy Toolbox (33) (v2.1; RRID:SCR_013273).

First, we cross-correlated the time courses of the voxels in one ROI with the time courses of the voxels in the ipsilateral other ROI resulting in separate Pearson correlation matrices for the for the left (314 AMY voxels 512 aHC voxels) and for the right hemisphere (361 AMY voxels. 564 aHC voxels) for each speech period (n=60) and participant (n=25) separately. For each correlation, we then averaged across all speech periods within one condition (n=10) resulting in one correlation matrix per hemisphere (n=2), participant (n=25) and condition (n=6). Next, we Fisher-z transformed the r values to z scores and applied a noise filter, excluding all values below the 33rd percentile and negative correlations. To compare the different conditions, we then calculated difference scores by subtracting the corresponding correlation matrices. For a group-level analysis, we averaged across participants, resulting in one correlation matrix per hemisphere and comparison.

To test for significance, we performed a permutation test where we shuffled all z scores 2000 times across both correlation matrices for each condition and each participant. Analogous to the observed values, we then calculated the difference scores between conditions, and averaged across participants for each permutation separately, resulting in 2000 correlation matrices per hemisphere and comparison. Next, we evaluated all observed z values with a cumulative distribution function (CDF) of the normal distribution (with the M and SD of the corresponding permutation test-based null distribution). As CDFs represent the probability for a random variable to be smaller or equal to any specific test value, we subtracted that probability from 1 to gain a meaningful p-value (one-sided). Because the resulting p-value distribution was nonparametric, we calculated the negative log10. On the negative log10 scale, a value of 1.3 corresponds to a p-value of 0.05 and thus 1.3 served as our significance threshold. Finally, we added the number of significant correlations from one voxel to all the voxels in the other ROI resulting in connectivity intensity maps, showing higher values for voxels with more interconnectedness to the other ROI. For visualization purposes, we spatially smoothed the connectivity intensity maps with a 2-mm full-width at half-maximum isotropic Gaussian kernel.

### GCA to estimate directional functional brain connectivity in AMY-aHC-AC network

To estimate the directional functional connectivity of the AMY, aHC, and other ROIs that were activated during the experiment, we conducted a Granger causality analysis (GCA). This analysis determines how the time course in one brain region can explain the time course in another brain area. If this determination is significant, we assume that one region is directionally connected to other region. GCA was performed with the MVGC Multivariate Granger Causality Matlab toolbox (34) (mvgc_v1.0; RRID:SCR_015755).

We defined the ROIs based on activations resulting from group-level contrasts between the feedback-based adaptive and non-adaptive conditions across and within the aggressive and joyful speech conditions. We thus included 10 ROIs in total from the bilateral AC (Heschl’s gyrus, mid- and posterior STC) and bilateral AMY and aHC. The exact MNI coordinates of the peak voxels of each ROI are marked with an asterisk in Supplementary Table S1. Cortical ROIs comprised voxels within a 3mm sphere around the peak location. For the AMY and aHC we used the same masks as created for the main experiment.

From each ROI, we extracted the time course of activity as the first eigenvariate across voxels within the ROI. For each participant, we epoched the ROI time course data for each trial in a time window from TR 8-44. We pooled all epochs across all participants instead of estimating the GCA for each participant separately. This is in accordance with previous fMRI studies (12, 13), since the number of trials for each participant is usually too low to provide an accurate estimation of brain connectivity. To mitigate this limitation, we additionally introduced a very conservative significance threshold to reduce the possibility for false positive results. We used F-tests to check for significance of the connections at p<0.01, FDR corrected. For each of the six experimental conditions, we ran a separate GCA. To compare the speaking modes, we subtracted the GCA results of the different conditions from each other for each emotion separately and applied the same conservative significance threshold. Supplementary Table S2 provides a full overview of the connectivity parameters between all ROIs.

### Post-experimental perceptual ratings and analysis

After the main fMRI experiment, we asked listening participants to dynamically rate the affective speech intonations heard inside the scanner during the fMRI experiment. Participants were still unaware of the different speaking conditions. For visual and statistical comparison, they also rated neutral intonations that we recorded from each speaker after the main experiment. Listening participants performed the continuous ratings twice on each of the 45.6-s speech periods, with one rating along the dimension arousal (low to high arousing; level of emotional engagement and intensity of own emotional reaction to vocalizations) and one along the dimension valence (negative versus positive valence; level of perceived negativity or positivity of the vocalizations), as was previously done in our recent study on music listening (35). As in our previous study (35), we expected that arousal and valence ratings on affective speech could be rated accurately and independently. The participants listened to the speech recordings through high-quality headphones and rated the speech periods by continuously moving a joystick, that was presented on a computer screen, up and down (arousal, from 0 to 1) or left and right (valence, from -1 to +1). The starting point for each speech sample was always set to the midpoint of the scale. The ratings were sampled at a frequency of 16 Hz.

To test for differences between the conditions and find out if there was an effect of sex of the speaker, we first averaged the rating values over an interval from the 9.6s to 45.6s (TR = 8 to end of speech period). The means of each listener were then subjected to a two-way repeated measures ANOVA (rmANOVA) with the two within-subject factors condition (neutral, joyful OnLfeed, joyful OnRfeed, joyful Offfeed, aggressive OnLfeed, aggressive OnRfeed, aggressive Offfeed) and sex (male, female). For a measure of effect size, we calculated the eta squared (η^2^) which describes the part of variance explained by the respective factor. The two-sided post hoc tests were Bonferroni-corrected for multiple comparisons. P=0.05 was considered statistically significant for both the rmANOVA and the post hoc tests. We applied Mauchly’s test to determine if the assumption of compound asymmetry was met. Significance of Mauchly’s test indicates that sphericity is not met and the estimate of sphericity ε indicates which adjustment to use (ε<0.75 Greenhouse-Geisser, ε>0.75 Huynh-Feldt) to correct for violations of sphericity. For both rating dimensions, Mauchly’s test indicated that the assumption of sphericity had been violated (valence: *χ^2^* (90)=247.76, p=1.1 10-16; arousal: *χ^2^* (90)=292.80, p=2.7×10^-23^), therefore degrees of freedom were corrected using Greenhouse-Geisser estimates of sphericity (valence: ε=0.303; arousal: ε=0.200).

### Acoustic analysis of adaptive and non-adaptive affective speech

We analysed the live and recorded affective speech intonations from the main experiment and the neutral intonations from the speech pre-recordings by using the openSMILE toolbox (36) (2.3; RRID:SCR_021273) which extracts acoustic features that are central to the analysis of affective speech. We extracted frequency-related parameters (e.g., pitch, jitter, formants), energy/amplitude-related parameters (e.g., loudness, shimmer), spectral (balance) parameters (e.g., alpha ratio, HBI Hammarberg Index, MFCC mel-frequency cepstral coefficient, spectral flux), and temporal features (e.g., rate of loudness peaks, duration of voiced and unvoiced segments). For all parameters, the mean and standard variation were quantified, resulting in a total of 88 acoustic voice features. Supplementary Table S3 provides a full list of all included features. We extracted the time courses of features for the 45.6-s speech periods in time bins of 1.2s in correspondence with the sampling rate of brain signals during the fMRI experiment (TR=1.2s).

Across the 88 acoustic features extracted from each speech period, we calculated difference scores between the conditions (e.g., OnLfeed minus Offfeed) for each participant and each emotion separately. These difference scores were subjected to two different analyses. First, we tested for statistical differences across all participants by applying two-sided one-sample t-tests (difference score against 0) for each of the 88 features (p<0.0001, FDR corrected). Second, we performed a principal component analysis with varimax rotation on the acoustic difference score profile of each participant. This was done with the aim to identify major acoustic vocal profiles of speakers that influence the perceptual and brain data of listeners.

### Correlation analyses between speaker and listener data

The time courses of vocal features were subjected to correlation analyses with the time courses of both perceptual rating dimensions and with the time courses of brain data in different regions (i.e., the same 10 ROIs as defined in the GCA). We downsampled the perceptual ratings to 0.8333Hz so that they would correspond to the sampling rate of the functional brain data (TR=1.2s) and acoustic features. To evaluate the relationship between vocal feature and brain activity, we shifted the vocal features by 3.6s (3*TR) to account for the delay of the BOLD signal (7).

We estimated the Pearson correlation coefficient of each acoustic feature to perceptual rating or brain data relationship for each single 45.6s speech period of the main experiment for each participant separately. This resulted in correlation coefficient distributions containing 250 coefficients for each acoustic feature within each of the six experimental conditions across all listening subjects (25 subjects x 10 speech periods per condition). For each of the six experimental conditions, the vocal feature–brain data relationships resulted in 880 distributions (88 vocal features x 10 ROIs) and the vocal feature–rating relationships resulted in 176 distributions (88 vocal features x 2 rating dimensions).

The correlation coefficients were transformed by using the Fisher-z transformation procedure, and the distributions of these transformed coefficients were fitted with a normal distribution, resulting in two fit parameters (M, SD). To test whether the distribution of correlation coefficients was significantly different from a normal distribution with M=0, we performed a permutation test to acquire a null distribution with an estimated mean and standard deviation. For the permutation, we shuffled the acoustic features (n=88) to speech period (n=10) relationship 500 times for each condition and each participant, resulting in 101,376 permutations for the entire sample. Each fitted distribution of the empirical data (empirical M and SD) was compared with the normal distribution resulting from the permutation sampling (permutation M and SD) by using two-sided Kolmogorov-Smirnov tests; significance was tested for each of the 88 acoustic features in each condition by using an FDR-corrected level of p<0.001. The same procedure as described above was also applied to the relationships between the time courses of the perceptual ratings and the brain data in all 10 ROIs, resulting in 20 correlation coefficient distributions (2 rating dimensions x 10 ROIs).

## RESULTS

### Amygdala-hippocampal connectivity is increased by adaptive affective speech

In the experimental setup, speakers were presented with a continuously changing vertical bar graph on a screen that represented the AMY-aHC connectivity feedback from listeners’ brains in real time. Based on this feedback signal, speakers optimized their affective speech expressions to maximize AMY-aHC connectivity. Instead of the entire HC, we quantified signal and connectivity only the anterior part of the HC (aHC) that is classified as the emotional subpart of the HC compared to the cognitive subpart in the posterior HC (12).

All speakers producing affective speech were trained actors who had extensive vocal training and were adept at introducing vocal variation by modulating their voices along various vocal intonation features. While quantifying the brain activity and especially the AMY-aHC connectivity using real-time functional magnetic resonance imaging (fMRI), the listeners were instructed to listen passively but attentively to the affective speech produced by speakers expressed in two socially relevant emotions (joyful and aggressive).

The affective speech was produced in three different speaking modes. In two feedback-based adaptive speech conditions, speakers adapted their speech in response to either the left or the right AMY-aHC connectivity, which we refer to as the OnLfeed and the OnRfeed conditions (together referred to as Onfeed). In the non-adaptive speech condition, speakers received no information about the AMY-aHC connectivity, which we refer to as the Offfeed condition. To determine if our main experimental manipulation for feedback-based adaptive speech was successful, we compared the AMY-aHC interconnectivity during the adaptive and non-adaptive speech conditions. We analyzed the AMY-aHC connectivity by performing correlation analyses between the ipsilateral AMY and aHC on a voxel-level. Specifically, the time courses of all voxels in one region-of-interest (ROI) were individually correlated with the time courses of all voxels in the other ROI. For contrasting the experimental conditions (e.g., [OnLfeed – Offfeed]), we then subtracted the Fisher z-transformed values of the correlations between conditions and determined the significance of each connection based on that difference score with permutation statistics. By then adding the number of significant correlations from one voxel to each of all the voxels in the other ROI, we obtained connectivity intensity maps showing higher values for voxels with more interconnectedness to the other ROI for each contrast. Because the AMY and aHC consist of several subregions, we determined the localization of the resulting connectivity patterns with probabilistic cytoarchitectonic maps (37). For visualization, we chose representative slices in the sagittal, coronal, and axial views whose approximate positions are illustrated in Fig. 2A.

Comparing AMY-aHC connectivity during feedback-based adaptive and non-adaptive speech ([OnLfeed – Offfeed] and [OnRfeed – Offfeed]), we found significantly increased AMY-aHC connectivity during adaptive speech, indicating a successful implementation of our experimental manipulation. Additionally, some more detailed observations could be made across both contrasts and emotions (Fig. 2B). First, the right-hemispheric AMY-aHC axis showed much higher interconnectivity compared to the left hemisphere, and the magnitude of connectivity in the right hemisphere varied between emotions. Second, left-hemispheric AMY-aHC connectivity was relatively lower and seemed to show little variance across emotions.

Third, joyful speech overall seemed to elicit higher AMY-aHC connectivity than aggressive speech, especially in the right hemisphere. Fourth, significantly connected AMY subregions were distributed across large parts of the structure with most voxels showing some degree of connectivity to the aHC. These areas with increased connectivity were located bilaterally in the laterobasal amygdala nucleus (LB) for aggressive speech and in the LB and superficial amygdala nucleus (SF) for joyful speech. Fifth, significantly connected aHC subregions were organized in distinctive clusters with discrete peak locations with a very high degree of connectivity to the AMY. Parts of the aHC located outside the high-connectivity clusters showed no or only a very low degree of connectivity to the AMY. Notably, the locations of the high-connectivity clusters were similar across hemispheres, affective tones, and contrasts. One cluster was located in the most anterior part of the aHC, close to or directly bordering the AMY, and extended medially (presumably belonging to the hippocampal-amygdaloid transition area (HATA) and the hippocampal area CA1). The second cluster was located more posteriorly in the aHC and extended more laterally, presumably belonging to either the subiculum or CA1. However, while the localization was bilaterally similar, clusters in the right hemisphere were much more extended and generally increased in connectivity with much higher peak values.

To evaluate if the two feedback-based adaptive speech conditions were equally effective in increasing AMY-aHC connectivity, we contrasted them against each other ([OnLfeed – OnRfeed] and [OnRfeed – OnLfeed]) (Fig. 2C). Across emotions, OnRfeed affective speech seemed to elicit slightly higher connectivity than OnLfeed speech in the right hemisphere, yet the difference was rather low. In the left hemisphere, both feedback-based adaptive speech conditions seemed to elicit almost the same degree of connectivity (Fig. 2C). Comparing the non-adaptive to feedback-based adaptive speech conditions showed similar spatial patterns but largely attenuated effects (Fig. S1).

Taken together, the highest interregional connectivity overall was found during feedback-based adaptive joyful speech (OnRfeed) in the right hemisphere in the LB and SF nuclei of the AMY, the medial anterior aHC (HATA and CA1), and the lateral posterior aHC (CA1 or subiculum).

### Adaptive affective speech drives neural activity in the auditory cortex

To evaluate the influence of the dynamic and feedback-based adaptive affective speech beyond the AMY-aHC connectivity level, we quantified the neural activity in listeners during the speakers’ adaptive and non-adaptive affective speech performance. Specifically, we were interested in neural effects in the core and extended voice processing network, especially in the AC, IFG, AMY, and aHC (3, 7). In the AC, we defined the cortical voice areas (VA) by a functional voice localizer scan (Fig. S2). The resulting VA cluster spread over large portions of the bilateral AC, with peaks located in the middle part of the higher-order AC labeled as Te3 that largely extends over the superior temporal cortex (STC) (Fig. S2).

Comparing feedback-based adaptive against non-adaptive speech ([Onfeed > Offfeed]) across emotions revealed clusters of activity covering large areas of the bilateral AC located within the VA with activity peaks in the STC, planum temporale (PTe), and the low-level Heschl’s gyrus (HG) as well as in the middle temporal cortex (MTC) belonging to the associative auditory cortex (Fig. 3a). While we did not statistically test for lateralization effects, the effect was overall noticeably more pronounced in the right hemisphere (Tab. S1a).

**Figure 3.**
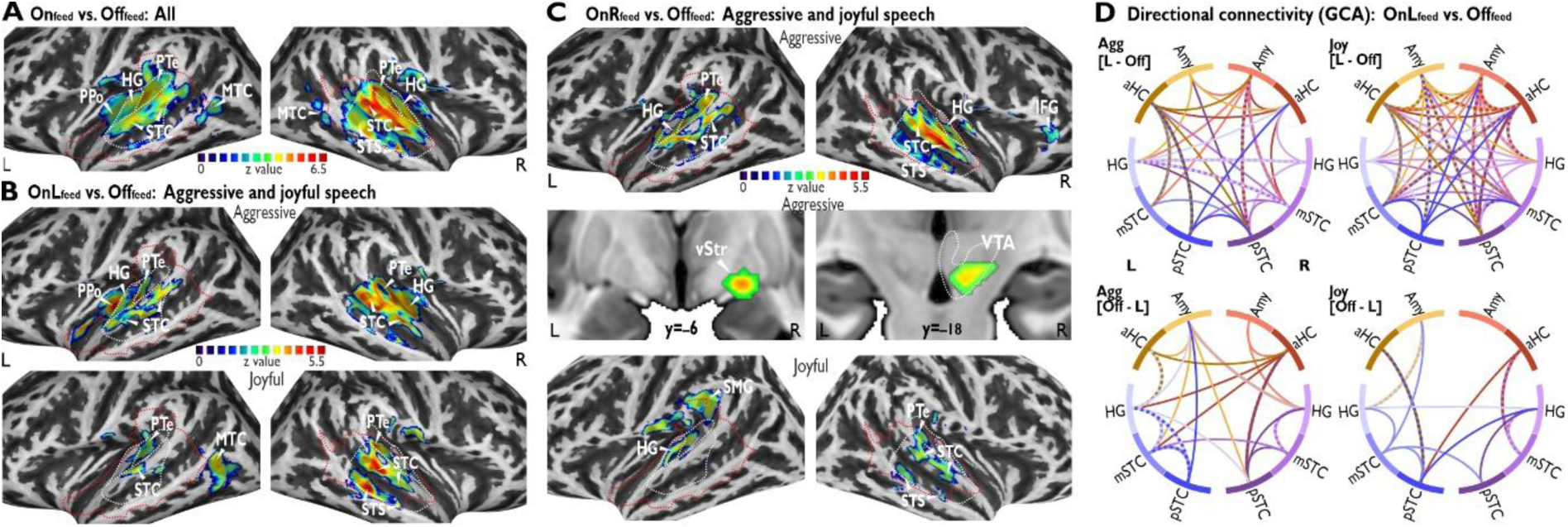
Adaptive affective speech drives neural activity and directed connectivity. (A) Neural activations for Onfeed versus Offfeed speech across both vocal affective tones, (B) for OnLfeed versus Offfeed for aggressive and joyful speech separately, and (C) OnRfeed versus Offfeed for aggressive and joyful speech separately. All contrasts showed only cortical activations except in C the middle panel shows also subcortical activations in vStr and VTA. The red dashed line delineates the VA and white dashed lines the anatomical subregions of the AC (as defined in Fig. 3). All neural activations thresholded at p<0.005, cluster size k>64 voxels, p=0.05 corrected on the cluster-level. See also Supplementary Fig. S2. (D) Functional connectivity determined by Granger causality analysis (GCA; p<0.01, FDR corrected) between the AMY, aHC, and lower- and higher-level AC (HG and m/pSTC, respectively) during aggressive (left column) and joyful speech (right column) for contrasting OnLfeed and Offfeed speech ([OnLfeed > Offfeed] in the upper row and [Offfeed > OnLfeed] in the lower row). Color of connections is defined by the region of origin; two-colored connections indicate bidirectional connectivity. See also Supplementary Fig. S2. Abbreviations: AC auditory cortex, VA voice area, AMY amygdala, aHC anterior hippocampus, HG Heschl’s gyrus, PTe/PPo planum temporale/polare, STC superior temporal cortex, STS superior temporal sulcus, IFG inferior frontal gyrus, MTC middle temporal cortex, SMG supramarginal gyrus, vStr ventral striatum, VTA ventral tegmental area, L/R left/right hemisphere, Agg aggressive, Joy joyful.

When comparing feedback-based adaptive against non-adaptive speech separately for each adaptive speech condition ([OnLfeed > Offfeed] and [OnRfeed > Offfeed]) and emotion (Fig. 3b-c), neural activity overall largely followed the above-mentioned pattern by mostly being restricted to the AC and being more pronounced in the right hemisphere. In aggressive speech, both OnLfeed and OnRfeed affective speech elicited highly similar AC activity with peaks in the bilateral STC, HG, and PTe (Fig. 3b-c, upper row). However, during OnRfeed speech we found additional smaller cortical and subcortical clusters. One cluster had peaks in the right IFG (pars triangularis and orbitalis), and one had peaks in the right ventral pallidum as part of the ventral striatum (vStr) and ventral tegmental area (VTA) (Fig. 3c, upper and middle row). In joyful speech, both OnLfeed and OnRfeed speech showed spatially greatly reduced left AC activity. However, we found additional activity in the left MTC during OnLfeed speech and in the left supramarginal gyrus during OnRfeed speech (Fig. 3b-c, lower left panels). In the right hemisphere, activity during both OnLfeed and OnRfeed joyful speech was not as pronounced in the HG but more focused on the STC and STS. Additionally, right-hemispheric activity during OnRfeed speech was less pronounced overall (Fig. 3b-c, lower right panels).

The reverse contrasts ([Offfeed > OnLfeed] and [Offfeed > OnRfeed]) yielded much less and much smaller activation clusters and in completely different and largely unrelated areas. Most importantly, we found no increased activity in the AC for both emotions (Fig. S3a-b). Offfeed aggressive speech showed increased occipital activation and Offfeed joyful speech showed increased activation in the prefrontal cortex (PFC) and thalamic areas.

Finally, we also compared both feedback-based adaptive speech conditions against each other ([OnLfeed > OnRfeed] and [OnRfeed > OnLfeed]). During OnRfeed speech, various areas showed higher activation than during OnLfeed speech (Fig. S3c). While both emotions elicited activity in the left dorsolateral PFC, OnRfeed aggressive speech showed additional frontal activity clusters with peaks in the bilateral dorsomedial PFC, left frontal operculum, and in the right IFG (pars orbitalis). OnRfeed joyful speech showed additional activations in the right dorsal striatum and the bilateral thalamus. The thalamus clusters were located in the left “auditory” thalamus (medial geniculate body) that is involved in more basic acoustic processing (3) and the right “limbic” thalamus (anterior nucleus) that delivers information to mainly limbic structures, especially also the HC (38). When comparing OnLfeed with OnRfeed speech only aggressive speech elicited higher activations, namely in one small cluster in the right precuneus (Fig. S3d).

Notably, both the left dorsolateral PFC and the thalamus clusters reported during joyful speech for the comparisons of [OnRfeed > OnLfeed] (Fig. S3c) as well as [Offfeed > OnLfeed] (Fig. S3a) were largely overlapping. This indicates that these activations were neither specific to OnRfeed nor to Offfeed speech. Additionally, OnRfeed aggressive speech showed increased activity in the right IFG for [OnRfeed > OnLfeed] (Fig. S3c) and [OnRfeed > Offfeed] (Fig. 3c). Since both IFG clusters were overlapping, this indicates that the right IFG activation is specific to OnRfeed aggressive speech. Furthermore, OnRfeed aggressive speech showed increased activity in a cluster with peaks in the vStr and VTA for [OnRfeed > Offfeed] (Fig. 4C). However, these areas did not show up for [OnRfeed > OnLfeed] (Fig. S2C) nor for [OnLfeed > Offfeed] (Fig. 3b). This suggests that the activations in the vStr and VTA were also slightly increased during OnLfeed aggressive speech but not as much as during OnRfeed and thus the activations did not reach significance [OnLfeed > Offfeed].

**Figure 4.**
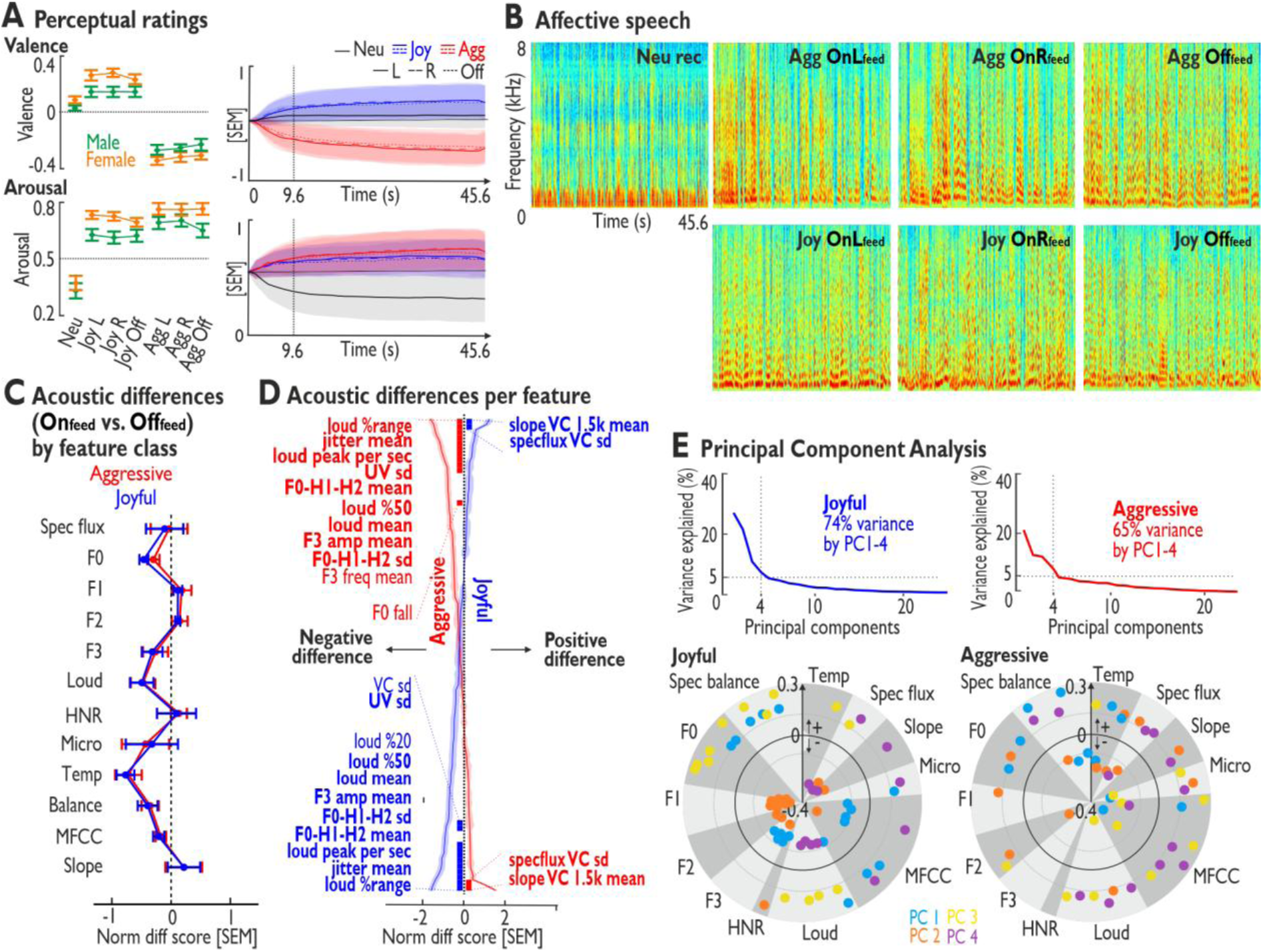
Analyses of listeners’ perceptual ratings and speakers’ acoustic data. (A) Mean (+1 SEM) perceptual arousal and valence ratings averaged over the 9.6–45.6s vocalization periods (left panel; male and female voices separated) from continuous dynamic ratings (right panels). See also Supplementary Fig. S3. (B) Spectrograms of example vocalizations for neutral voices and for aggressive and joyful affective adaptive speech (OnLfeed and OnRfeed) and non-adaptive speech (Offfeed). (C) Normalized difference scores for 88 acoustic vocal features (Onfeed minus Offfeed) summarized by acoustic feature class. See also Supplementary Fig. S4. (D) Normalized difference scores for each of the 88 vocal features. Features are sorted from maximum positive to maximum negative differences (joy) or vice versa (aggression); thick bars indicate significant differences (two-sided one-sample t-tests, p<0.0001, FDR corrected) with the respective features listed alongside, and a bold print indicates corresponding features for both emotions. (E) Principal component analysis (PCA) across acoustic difference scores for aggressive and joyful speech. Scree plots for both emotions (upper panel) and profile plots of factor loadings for PCs 1–4 for joy and aggression sorted by major acoustic feature classes (lower panel); only loadings above +/- 1.5 SD are shown. See also Supplementary Fig. S5 for more detailed profile plots. Abbreviations: agg aggressive, joy joyful, neu neutral, Spec flux spectral flux, F0 fundamental frequency, F1–F3 formants 1–3, Loud loudness, HNR harmonics-to-noise ratio, Micro micro irregularities, Temp temporal, Balance spectral balance, MFCC Mel-frequency cepstral coefficients, Slope spectral slope, VC/UV voiced/unvoiced segments, F0-H1-H2 harmonic difference, amp amplitude, freq frequency, PC principal component.

Taken together, only feedback-based adaptive affective speech elicited increased activity in the bilateral AC and especially in the right AC. Additionally, adaptive aggressive speech elicited more AC activity than joyful speech and OnRfeed aggressive speech additionally induced right IFG and subcortical activity.

### Higher interconnectivity in auditory-limbic brain network during adaptive affective speech

As the brain of listeners showed differences in AMY-aHC connectivity and regional neural activity between feedback-based adaptive and non-adaptive speech, we also aimed to explore if adaptive speech had any influence on the connectivity in a broader neural network including the AMY and aHC and central areas of the voice processing network in the AC. To examine the bilateral AMY-aHC-AC network, we determined directional connectivity using a Granger causality analysis (GCA), separately for joyful and aggressive speech (Fig. 3d). To define the ROIs for the GCA, we chose regions that were active throughout all contrasts of the feedback-based adaptive versus non-adaptive speech conditions, thus limiting us to regions in the bilateral AC (mid- and posterior STC, HG; see Supplementary Table S1 for the MNI coordinates of the peak voxels of each ROI). Additionally, we entered the bilateral AMY and aHC as ROIs as they represented the main experimental focus with the same anatomical masks as used during the fMRI experiment (Fig. 2a).

When comparing the AMY-aHC-AC network during feedback-based adaptive and non-adaptive speech, we found increased interconnectivity between all our ROIs for both emotions (F-statistics, p<0.01, false discovery rate (FDR) corrected) (Fig. 3d for OnLfeed versus Offfeed; Fig. S3e for OnRfeed versus Offfeed). This was reflected in a higher number of connections overall as well as a higher percentage of interhemispheric connections between all ROIs. Additionally, the AMY and aHC were much more involved in the network during Onfeed speech, both in total number of connections and proportionally to the whole AMY-aHC-AH network for incoming as well as outgoing connections. While both emotions showed higher interconnectivity and higher AMY and aHC involvement in Onfeed than Offfeed speech, the difference was much more pronounced during joyful speech. Also, during Onfeed joyful speech the AMY and aHC were interconnected within both hemispheres (Fig. 3d; Fig. S3e, upper right panel). During Onfeed aggressive speech, the right aHC and AMY seemed to be also connected, this was however not the case in the left hemisphere (Fig. 3d; Fig. S3e, upper left panel). Nevertheless, both limbic regions were highly interconnected with the AC areas and also showed connectivity with the contralateral AMY and aHC.

While both Onfeed speech conditions showed highly similar connectivity patterns in the corresponding emotions when compared to Offfeed speech, overall, the increase in interconnectivity for Onfeed speech seemed to be more pronounced during OnLfeed for joyful speech (Fig. S2f, right column) and during OnRfeed for aggressive speech (Fig. S3f, left column). This was true for both, the total number of connections in the whole network as well as when only regarding the AMY and aHC. Supplementary Table S2 provides a full overview of the connectivity parameters between all ROIs.

### Perception of the emotional intensity and quality of affective speech

To explore if listeners recognized the affective tone of the vocalizations and if the perception differed between feedback-based adaptive and non-adaptive speech, we also assessed different dimensions of perceptual ratings of all speech intonations. Listeners rated all vocalization samples they listened to during the main experiment on emotional intensity (i.e. arousal; how intensely do the listeners perceive the speech? How engaged are they emotionally while listening?) and affective quality (i.e. valence; how negatively or positively do the listeners perceive the speech?) (Fig. S4). All speech periods were cut to 45.6s, which is the duration of each trial during the experiment. We quantified the mean ratings from 9.6-45.6s (Fig. 4a), which was the vocalization interval also used during other analyses (see Methods section, Online functional brain data analysis). In addition to the affective speech samples, we included neutral speech as a reference condition recorded by each speaker after the main experiment.

Valence ratings differed between affective speech conditions (rmANOVA, p<0.01, n=25, Greenhouse-Geisser (GG) adjusted p-values, main effect condition: F6,144=79.53, p=3.16 10-18, η^2^=0.631). Listeners perceived aggressive speech as more negative than neutral speech (posthoc test, p<0.01, FWER corrected, all ps<5.42x^10-7^). For joyful speech, the comparison did not reach significance (all ps>0.088), but it was rated much more positive than aggressive speech (all ps<7.28x^10-8^). Speech during all experimental conditions, including Onfeed as well as Offfeed speech periods, was rated as having a similar emotional quality (all ps>0.326). Within the experimental conditions, intonations from male and female speakers were rated slightly different (interaction of condition*sex: F6,144=9.20, p=6.72x^10-6^, 2=0.022). Female speech seemed to be rated as more intense in their emotional quality than male speech. However, this difference only reached significance for joyful speech during both Onfeed conditions (ps<0.002). Additionally, OnRfeed joyful speech produced by female speakers was perceived as more positive than neutral speech (p=0.001).

Arousal ratings for affective speech were overall different from neutral speech (rmANOVA, n=25, GG-adjusted p-values, main effect condition: F6,144=32.92, p=1.37x^10-8^, η^2^=0.382) with any affective speech being more arousing than neutral speech (all ps<3.9x^10-5^). Joyful and aggressive speech were perceived as equally arousing (all ps>0.055) and Onfeed and Offfeed speech did also not differ in perceived emotional intensity (all ps>0.620). Additionally, speech expressed by female speakers was overall perceived as more arousing than speech expressed by male speakers (main effect sex: F6,144=37.36, p=2.59x^10-6^, η^2^=0.040).

### Acoustic properties of adaptive affective speech differ from non-adaptive speech

As the different affective speech conditions seemed to elicit different neural activity and connectivity patterns, we investigated if the speakers’ affective speech samples also differed in their acoustic properties (Fig. 4b). Therefore, we quantified 88 central vocal features commonly found in affective speech communication (39) in terms of vocal pitch, formant properties, loudness, and other spectral and temporal characteristics (Fig. S6 and Tab. S3 for a full list of features). To compare the acoustic properties between the different speaking conditions, we calculated difference scores (e.g. Onfeed minus Offfeed) on which we based all following analyses.

When comparing feedback-based adaptive and non-adaptive speech ([OnLfeed – Offfeed] and [OnRfeed – Offfeed]), the difference profiles over the broad acoustic feature classes for both contrasts were almost identical (Fig. S5a-b). Additionally, a direct comparison between both feedback-based adaptive speech conditions showed very little difference over the broad acoustic feature classes (Fig. S5c, left panel). Moreover, in a more detailed analysis comparing each feature individually (two-sided one-sample t-tests, difference score between OnLfeed and OnRfeed against zero, n=25, p<0.0001, FDR corrected), none of the feature differences reached significance (Fig. S5c, right panel), indicating highly similar acoustic properties in both feedback-based adaptive speech conditions. Thus, we analyzed the acoustic differences between feedback-based adaptive and non-adaptive speech across the two adaptive speech conditions (i.e., [Onfeed – Offfeed]).

For feedback-based adaptive versus non-adaptive speech, the majority of features showed negative values in terms of lower values in these features in adaptive speech. Additionally, features in joyful and aggressive speech seemed to be highly similar. Over the broad acoustic feature classes, the biggest differences between conditions in terms of lower values during Onfeed speech seemed to be related to temporal parameters, followed by features related to loudness, pitch, spectral balance, and the third voice formant (F3) (Fig. 4c). A more detailed analysis of the individual features (two-sided one-sample t-tests, difference score between Onfeed and Offfeed against zero, n=25, p<0.0001, FDR corrected) showed that Onfeed speech was higher in features related to spectral voice variations (mean of the spectral slope and variation of the spectral flux) for both affective tones (Fig. 4d). Lower feature values in Onfeed speech were found in parameters related to loudness (mean, range, 50th percentile), temporal characteristics (loudness peaks per second, variation in the duration of unvoiced (UV) segments), mean jitter, spectral features (mean and variation of the harmonic difference H1-H2; F0-H1-H2), and mean F3 amplitude (Fig. 4d). Onfeed joyful speech additionally showed lower variation in the duration of voiced (VC) segments and a lower 20th percentile of loudness. Onfeed aggressive speech was additionally characterized by lower mean F3 frequency and a flatter slope of falling pitch (less melodic intonation) than Offfeed speech (Fig. 4d).

As listeners might be differently sensitive to various features and since it was the aim of the speakers to individually adapt their voices to the listeners’ reaction, we wanted to explore if there were different vocal profiles across speakers within each affective tone. Therefore, we performed a principal component analysis on the acoustic difference profiles ([Onfeed – Offfeed]) for the 88 features of each listener and affective tone (PCA; n=25; Fig. 5E; Fig. S6 for a more detailed view). Thereby, we could identify four principal vocal profiles separately for aggressive and joyful speech, with each acoustic profile explaining >5% of the overall variance and all of them together explaining 74% variance for joyful and 65% variance for aggressive speech.

**Figure 5.**
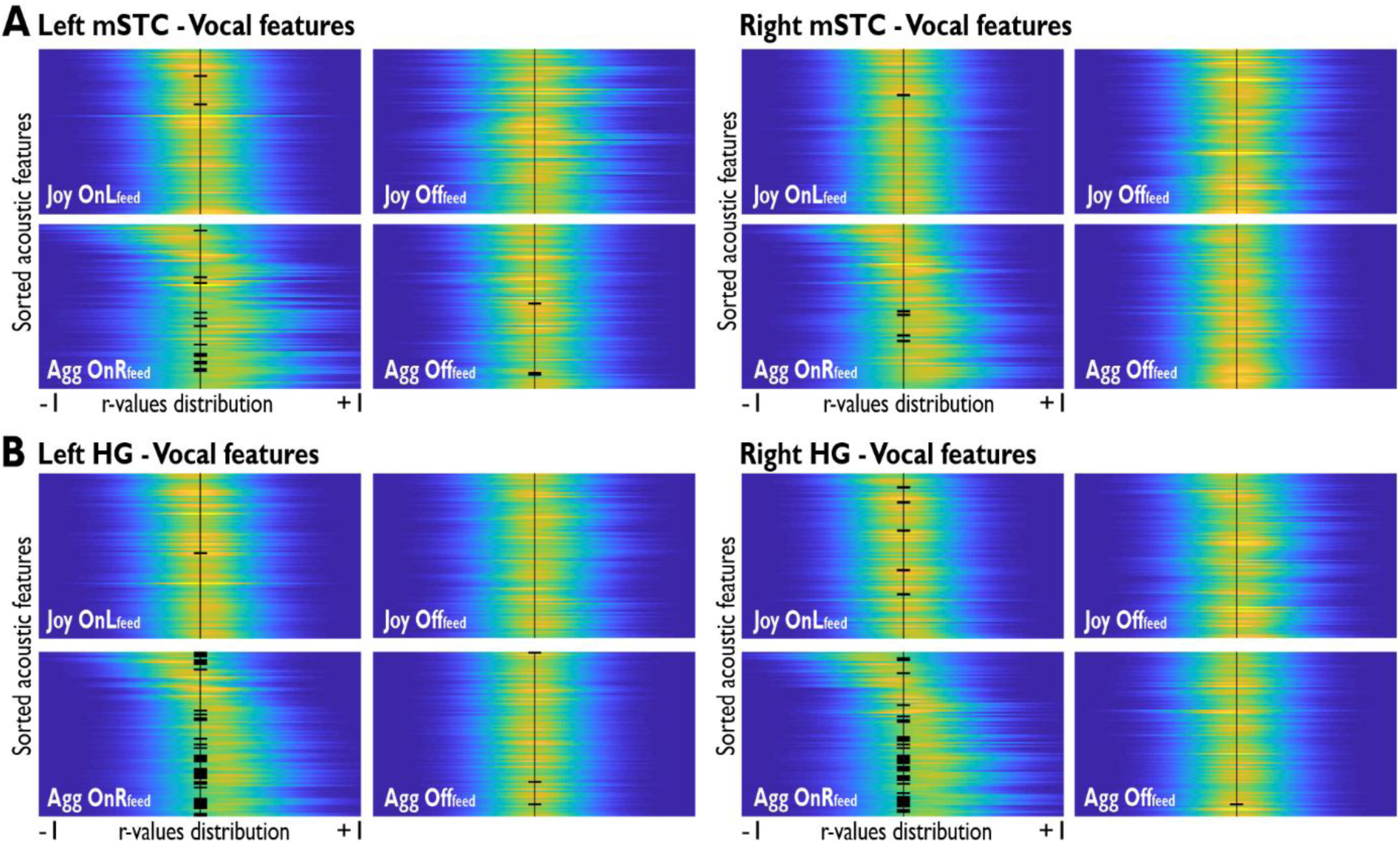
Communicative coupling of speakers’ voice features with listeners’ neural AC dynamics. (A) Density distributions (x-axis; yellow/high density, blue/low density) of Pearson correlation coefficients (each line/distribution n=250, 25 subjects x 10 trials) between the 88 acoustic vocal features (y-axis) and neural activity in the left and right mid-superior temporal cortex (mSTC) during aggressive (agg) and joyful (joy) speech. (B) Density distributions for the left and right Heschl’s gyrus (HG). The features for joyful speech are sorted according to the mean of the distribution during the OnLfeed condition from negative to positive values; the same sorting order was applied to the Offfeed condition. For aggressive speech, the same sorting procedure was applied but according to the mean of the distribution during the OnRfeed condition. Black thick bars in the middle of the plot show a significant difference of the distribution from the null distribution (Kolmogorov-Smirnov test, p<0.001, FDR corrected). See also Supplementary Fig. S6 and S7 for more detailed plots.

The first principal acoustic profile for aggressive speech showed differential loadings on features related to pitch, spectral balance, and temporal parameters. The second principal acoustic profile showed differential loadings on features related to pitch, formants F1–F2, micro irregularities, spectral slope, spectral flux, spectral balance, and temporal parameters. Thus, we termed the first component a “dynamic pitch” vocal profile and the second component a “changing tone” vocal profile. Analogous with their major feature loadings we termed the third component “loud timbre” and the fourth component “loud spectral” vocal profile. For joyful speech, the profiles were a bit different. The first component showed loadings on features related to pitch, formant F3, spectral balance, and vocal timbre (Mel-frequency cepstral coefficient; MFCC); and the second component showed high loadings on harmonics-to-noise ratio (HNR) and negative loadings in features related to formants F1–F3, spectral flux, and spectral slope. Based on the different patterns of factor loadings, we thus termed the four profiles “harmonic timbre”, “dynamic tone”, “powerful pitch”, and “varying loudness” for joyful speech.

### Communicative coupling of speakers’ voice features with listeners’ neural AC dynamics

As speakers modulated their voices in adaption to listeners’ AMY-aHC connectivity, we hypothesized that there should be a coupling of speakers’ acoustic and listeners’ neural parameters. Therefore, we performed correlation analyses between the speakers’ acoustic and the listeners’ neural data. The analysis resulted in empirical correlation coefficient distributions, which were compared with a null distribution determined by permutation sampling using two-sided Kolmogorov-Smirnov tests (p<0.001, FDR corrected) (Fig. 5; Fig. S7-8). We correlated the 88 acoustic features quantified in the previous step with neural activity in the same regions as during the functional connectivity analysis, namely in the AC the high-level mid- and posterior STC and the low-level HG and subcortically the AMY and aHC.

While we found no correlations with the regional activity in the AMY and aHC, the bilateral AC (mSTC, HG) showed coupling with multiple acoustic parameters of the speakers’ affective intonations (Fig. 5). Overall, these acoustic-brain couplings revealed five key features. First, Onfeed speech showed a higher number of significant correlations. Moreover, during Offfeed speech the acoustic-brain dynamics only rarely correlated at all. Second, aggressive speech showed a much higher number of significant correlations than joyful speech. Third, neural activity in the HG (Fig. 5b) showed overall more coupling with speech features than activity in the mSTC (Fig. 5a). Fourth, in the mSTC we found more correlations in the left hemisphere, while in the HG the number of correlations was bilaterally equal. These latter two findings are mainly applicable to aggressive speech, as coupling during joyful speech was too low to reliably make these observations. Lastly, regarding the Onfeed conditions, we revealed differences between affective tones. During joyful speech, only OnLfeed speech showed significant correlations. Contrarily, during aggressive speech only OnRfeed speech showed significant correlations. This was true for the mSTC and HG in both hemispheres (Fig. 5).

When exploring the couplings on a more detailed level, we observed primarily positive and only a few negative acoustic feature-brain activity correlations. During OnLfeed joyful speech, the left mSTC signal correlated positively with the mean frequency of formant F3 and negatively with the mean of vocal timbre-related MFCC1 (Fig. S7a). For the right mSTC we found only a positive correlation with pitch height (Fig. S7b). In the bilateral HG, activity was positively associated with the mean F3 bandwidth (Fig. S8). For the right HG, we found additional positive correlations with the mean F2 frequency and loudness peaks per second, and negative correlations with the variation in F1 bandwidth and mean shimmer (Fig. S8b).

During OnRfeed aggressive speech, we found many more and predominantly positive correlations, especially related to formant properties and variation in multiple acoustic characteristics. The left mSTC signal was positively associated with parameters related to mean levels of formants properties (F1 frequency, F1–F3 bandwidth), pitch height, vocal timbre (MFCC3), and HNR. Additionally, we found positive correlations with acoustic features related to variation in spectral balance (Hammarberg index; HBI), spectral flux, and loudness. Lastly, loudness peaks per second correlated negatively (Fig. S7a). The right mSTC showed the same positive correlations as the left mSTC with those features that quantify variation (i.e., variation in HBI, spectral flux, and loudness) and additionally with mean shimmer (Fig. S7b). Regarding the HG, we found a lot of correlations with the same features as in the mSTC and additionally with many more formant-related and spectral features (Fig. S8). Specifically, bilateral HG activity correlated positively with features related to mean levels of formant properties (F1–F3 frequency and bandwidth), pitch height, spectral slope, spectral balance (HBI), spectral flux, and loudness. Moreover, the HG signal correlated positively with the same variation-related features as in the bilateral mSTC (variation in spectral balance (HBI), spectral flux, and loudness) and additionally with variation in spectral slope and formant properties (F1–F3 frequency and F1–F2 bandwidth). Negative correlations were found with the spectral balance parameter alpha ratio (in voiced segments), the vocal timbre-related MFCC4, and loudness peaks per second. Additionally, HG activity in the left hemisphere was positively correlated with the harmonic difference H1-A3 and negatively correlated with the amplitude variation in F2–F3 and the mean duration of unvoiced periods (Fig. S8a). In the right HG we found additional positive correlations with the mean and variation of shimmer, additional MFCCs, and mean HNR (Fig. S8b).

## DISCUSSION

Using a closed-loop real-time neuroimaging setup, speakers could successfully induce higher connectivity in the AMY-aHC circuit during feedback-based adaptive compared to non-adaptive affective speech. This finding has important implications. First, speakers can directly target and actively influence AMY-aHC connectivity in listeners with affective speech and specific AMY-aHC interactions seem therefore indeed crucial for affective speech processing. Importantly, the use of a dynamic and individually adaptive manipulation condition and its comparison to a condition with no direct manipulation allows to explicitly assign the resulting neural dynamics to this experimental difference. This substantiates the significance of the so far neglected role of the AMY-aHC interaction in affective speech processing (40). Second, this highlights the importance of considering variabilities in sensitivity and affect decoding abilities across individuals in affective speech experiments (41). Previous studies with non-adaptive affective speech samples might accordingly not have captured the full processing capacities of the limbic system that decodes affect from speech (3). Our experimental design provides speakers with the possibility of adapting their affective speech to the individual emotional and neural sensitivity in listeners and thus potentially resembling real-life affective communication dynamics more closely. This might furthermore corroborate the notion that the decoding of affect from speech in the limbic system was evolutionarily shaped by and is thus maximally optimized during dynamic and social real-life interactions (42).

Apart from generally observing that adaptive and dynamic affective speech can directly be modulated to increase and potentially maximize AMY-aHC connectivity, we also found differential effects in connectivity strength and similarities in connectivity patterns of AMY and aHC subregions across conditions and affective tones. Regarding connectivity strength, a first important observation was that adaptive joyful speech seemed to increase the AMY-aHC connectivity more than adaptive aggressive speech. Joy compared to aggression is a more pro-social and interactive emotional expression (43) that however often depends on contextual associations (44). This emotion-context-association might be critically decoded by synchronization of the AMY-aHC circuit. Second, connectivity in the right AMY-aHC circuit was overall higher than in the left AMY-aHC circuit even for the condition of stimulating the left AMY-aHC circuit (OnLfeed). This might point to a predominance but not exclusivity of the right AMY-aHC circuit for affective speech deducing in line with a predominance of integrative versus analytic affective sound decoding (8) and affective versus cognitive theory of mind in right medial limbic systems (45) as well as a preference for melodic sound features (46) and emotional associative processes in the right brain (47, 48).

While the strength of connections differed between brain hemispheres and affective tones, the spatial distribution of the connectivity pattern in the AMY-aHC circuit was rather similar across conditions. These spatial distributions resemble known afferent and efferent connectivity patterns in the AMY-aHC circuit and thus they seemed functionally meaningful (8). Regarding the AMY as the seed region for outgoing connections, the LB nucleus was the main area with connections to the aHC during affective speech decoding. The LB nucleus is one of three main subnuclear groups of the AMY and is the main input region for auditory stimuli (11, 49) as well as the main input and output region for connections with the aHC (10, 13). Additionally, AMY voxels with increased connectivity to the aHC were spread among the majority of the LB complex. A recent study confirmed this pattern, with afferents from the AMY to the aHC being dispersed widely among the LB (10).

Regarding the aHC as the seed regions for outgoing connections, we revealed a spatially less distributed pattern including rather specifically circumscribed areas showing increased connectivity to the AMY. Specifically, large aHC sections showed only very low or no connectivity to the AMY. Regions with increased connectivity to the AMY were located in the subiculum, CA1, and HATA subregions of the aHC. All three of these subregions show high consistency with the well-established afferent and efferent connectivity of the aHC (8). The CA1 and subiculum are output nodes of the aHC and receive the majority of input from the LB complex of the AMY (10, 50). The HATA is the transitional area between the aHC and AMY and thus also involved in information exchange between these two areas (51). Additionally, the rather focused peak connectivity clusters seem to confirm that aHC efferents seem primarily organized in parallel circuits where each projection to a specific AMY subarea emerges from a discrete cluster of neurons that projects to only one efferent area (10).

Thus, the subregions with increased interconnectivity in the AMY-aHC circuit are anatomically and functionally reasonable (52), which specifically point to the relevance of the AMY-aHC connectivity in affective speech processing (8). We of course have to emphasize that the spatial resolution of functional magnetic resonance imaging is often too low to identify AMY and aHC subregions in all detail. But the observed increased AMY-aHC pattern together with the less pronounced connectivity pattern during non-adaptive affective speech supports the conclusion that speakers could indeed specifically target and maximize the AMY-aHC connectivity by individually adapting their affective intonations in real time.

Additional to a significant influence of adaptive affective speech on intralimbic circuits, the stimulation of AMY-aHC connectivity also evoked higher activity in a broader brain network. Adaptive speech across affective tones elicited much higher activation in the bilateral AC compared to non-adaptive affective speech. Regions in the AC are part of a central voice processing network that is also sensitive to affective voices (5–7, 53). This affective voice processing network typically also includes the AMY and aHC, which are structurally and functionally connected with the AC (8, 21). While the roles of both limbic regions and their functional interaction with the AC during affective sound and speech processing have been explored before (21, 54, 55), there have only been theoretical assumptions about the importance of the limbic AMY-aHC connectivity in the broader brain network. We provide empirical evidence to support this assumption, suggesting that synchronization of affect decoding processes in the limbic AMY-aHC circuit potentially leads to an increased information exchange of limbic circuits with auditory analysis processes in the AC. The AMY and aHC typically evaluate the relevance and associative meaning of the affective stimuli for the listener (3). The AMY is responsible for the affective evaluation and operates on a shorter timescale while the aHC provides a contextual frame and examines auditory information on a broader timescale (8). Specifically, the aHC supports temporal tracking and integration of affective voice information. This is especially important in the auditory domain as sounds usually develop over time especially in dynamic and adaptive speech (21, 56). This information seems relayed to the AC for sensory auditory analysis in low-order AC (HG, PTe) and for information integration in high-order AC (STC) (3).

While the activation patterns for adaptive compared to non-adaptive speech and for both affective tones were largely similar concerning the AC, they showed differences in other brain areas. For adaptive joyful voices, we found activity in the left higher-order AC (MTC) during OnLfeed speech and in the left supramarginal gyrus during OnRfeed speech. Both regions are relevant for decoding affective prosodic pattern with varying levels of temporal variations (3, 57). For adaptive aggressive voices, only OnRfeed speech showed additionally activated areas, namely in the right IFC, right vStr, and VTA. These activity clusters and their interaction with the AC have central evaluative functions in affective speech processing (3, 9, 57). The VTA is commonly assumed to decode pleasure and reward in sound (3, 58). But VTA activity was also found during aversive experiences (59, 60) and in salience detection in the absence of reward (61), which could both be very relevant during the perception of dynamic affective speech. When directly comparing the two adaptive speech conditions, almost exclusively OnRfeed speech induced increased activity compared to OnLfeed, namely in prefrontal and inferior fronto-insular areas relevant for affect classifications during both affective tones (21), and additionally in the thalamus and dorsal striatum during joyful voices to potentially track prosodic variations in vocal expression of joy (62).

As we specifically target the AMY-aHC connectivity in our experiment, an important observation was that we did not find increased activity in the AMY and aHC during adaptive speech. We have to note however that speakers aimed to increase the connectivity between the AMY and aHC rather than the absolute activity in both regions. The standard experimental approach based on the strength of limbic activity for non-modulatory (6, 63) and real-time neurofeedback neuroimaging studies (7) substantially differs from the approach in this experiment as we focused on the comparison of adaptive to non-adaptive affective speech in terms of their potential to stimulate neural connectivity.

Based on the increased intralimbic AMY-aHC connectivity and increased activity in low and higher-order AC areas during adaptive affect speech, we further examined neural dynamics in the broader cortico-limbic brain network for affective voice processing. Using a functional connectivity analysis, we again observed an increased interconnectivity between the AMY and aHC in adaptive versus non-adaptive speech for both affective tones. This AMY-aHC connectivity pattern based on averaged time courses across the voxels of each regions complements and confirms the previously described voxel-wise connectivity patterns. Additionally, the AMY-aHC circuits in both hemispheres were interconnected during adaptive joyful speech, which was only the case for the right AMY-aHC circuit in adaptive aggressive speech. This again potentially points to the enhanced emotion-context-association during the expression and perception of joyful speech. Apart from increased AMY-aHC connectivity, both limbic areas were also more interconnected with low and higher-order AC areas and showed a higher integration in the broader network during adaptive speech. This supports the suggestion that the synchronization of processes in the AMY and aHC during affect decoding increases the exchange of information with as well as the auditory and integrative analysis in the AC. When comparing the adaptive conditions with each other regarding the functional connectivity network, OnLfeed speech seemed to increase connectivity in the cortico-limbic network during joyful expressions and OnRfeed speech in turn increased cortico-limbic connectivity during aggressive expressions. This might point to an overall brain lateralization for processing positive and negative in the left and right human brain, respectively (64, 65). Although clear and consistent evidence for this valence hypothesis of brain lateralization effects for different emotions is so far missing (66), adaptive affective speech that aims to maximize affect decoding in the human brain might enhance this lateralization effect to some degree.

Extending the level of information about listeners’ responses beyond their neural signals and determining if the emotions were correctly induced, we also post-experimentally quantified subjective perceptual ratings of emotional intensity (arousal) and quality (valence) to the adaptive and non-adaptive affective speech intonations. Overall, aggressive speech was rated more negatively and more arousing than neutral speech, while joyful speech was perceived as more arousing but not as more positive than neutral speech. Yet, first, joyful speech was still rated as much more positive than aggressive speech and second, neutral speech can be perceived to have a positive quality but differing from joyful by having low arousal levels (39), as it is the case here. As adaptive and non-adaptive speech differed on the neural level, we also tested for perceptual differences, which we did not find. However, first, it is important to note that arousal and valence are just dimensions and thus a simplification of emotion perception. While they are useful in distinguishing aggressive and joyful speech (39), they might not be suited or sufficiently sensitive to capture the perceptual differences that arise from adaptive and non-adaptive affective speech, especially since the HC is also sensitive to more subtle modulations in arousal and valence (21). Second, it might be that the differences lie in the variation rather than the mean since this was the goal of the real-time setup.

Concerning the acoustic pattern of adaptive affective speech that seemed to enhance neural activity and neural connectivity patterns, affective speech in both adaptive conditions (OnLfeed, OnRfeed) was acoustically highly similar, but adaptive affective speech significantly differed from non-adaptive speech. First, this suggests that speakers did indeed vary their affective intonations between adaptive and non-adaptive speech. Second, speakers indeed seemed to adapt their speech according to listeners’ limbic connectivity feedback in specific features. Predominant features that were more pronounced in adaptive compared to non-adaptive speech were largely related to spectral voice variations for both affective tones, such as the spectral slope (steepness of the spectral profile) and spectral flux (spectral change over time). Thus, adaptive affective speech had a more distinct spectral profile and spectrally changed considerably over time. Unlike this enhanced spectral profile of adaptive affective speech, other features were rather suppressed in adaptive compared to non-adaptive speech, such as temporal micro irregularities (vocal jitter), formant and harmonics features (harmonics H1-H2, F3 voice formant) and loudness parameters (loudness peaks per second, loudness mean and range). This overall indicates more smooth, less hectic, and quieter affective vocalizations in the adaptive speech conditions. Given that the temporal integration windows for calculating the real-time AMY-aHC connectivity was rather extended, speakers might have introduced a distinct but consistent affective voice profile over time, which could lead to more authentic affective vocalizations (67). The lower harmonicity in adaptive speech could have further increased the authenticity of the vocalizations (68).

Certain acoustic features of affective speech thus seem to stimulate the intralimbic connectivity and neural activity in a broader brain network across all listeners. But adaptive affective speech is to a certain degree also listener-specific. We accordingly found four distinctive vocal profiles for each affective tone that indicate some level of listener specificity and speaker adaptivity during affective speech communication. These vocal profiles might reflect these different affective speaking styles, such as joyful amusement and elation (69) or aggressive anger and irritation (64), which could be strategically used and dynamically changed by speakers to achieve maximum emotional effects in listeners.

We therefore finally explored the coupling of speakers’ acoustic profiles with listeners’ neurocognitive parameters that might underlie the affective speech effects in the listeners’ brains. No acoustic-brain correlations were found in the AMY and aHC as those regions are usually not directly involved in affective voice features decoding (55) or only have a supplementary role for affective sound feature evaluations (6). But almost exclusively during adaptive speech, we could show coupling of acoustic features of affective speech produced by speakers with neural activity in the low AC (HG) and higher-order AC (mSTC) in the listeners’ brain. The HG showed more correlations overall than the mSTC and coupling in the HG seemed to be similarly pronounced in both hemispheres while coupling in the mSTC was higher in the left than in the right hemisphere. The HG is commonly bilaterally involved in decoding voice features that help to discriminate affective tones (70) and to detect important acoustic changes in affective sounds (35). The mSTC as part of the higher-order AC seems to show a lateralization in decoding acoustic features with the right hemisphere analyzing on a broader time scale to evaluate spectral properties (71, 72). Thus, the short-term changes in single acoustic features could be too specific to capture this more general process in the right but not in the left hemisphere. This notion is supported by the fact that the few features that did correlate with the neural activity in the right mSTC were almost exclusively related to features that quantify variation over time.

Besides these differential regional effects, we also found differential condition effects. Adaptive aggressive speech evoked much more correlations than adaptive joyful speech. Joyful voices potentially require more temporal information integration to be accurately recognized compared to aggressive voices (73). The acoustic features of joyful speech might thus not directly correlate with the neural signal but rather evoke a higher neural activation for representing integrated voice information (3) that are potentially less well captured by correlation analyses (35). Unlike for joyful speech, three important voice features correlated positively with both regions in both hemispheres for adaptive aggressive speech, namely variations over time in the spectral high-to-low frequency ratio, in spectral flux, and in loudness. The first two features are important parts of aggressive voice patterns (74) and they have strong perceptual effects in listeners for affective tone discrimination (54, 75). Furthermore, they can be easily modulated in adaptive affective speech (7) giving speakers a dedicated voice feature to affect emotional states in listeners. Regarding loudness, acoustic analysis of adaptive speech showed an overall lower mean and range than non-adaptive speech but no significant difference in loudness variation. Combined with the correlation results these acoustic findings encourage the conclusion that loudness was more subtly modulated than merely by increasing the loudness or changing between loudness extremes and that loudness variation is only influential when adapted to a listener’s individual reaction. The latter point also reinforces the argument that neural and acoustic differences between adaptive and non-adaptive speech were not simply a result of the task instruction that might have lead speakers to generally introduce more variance in their intonations during adaptive speech regardless of the listeners’ neural feedback, Only in adaptive speech did variation in specific features directly correlate with listeners’ neural activations which emphasizes the listener-specific adaption of the speakers’ affective intonations.

In summary, our data point to the effectiveness of adaptive affective speech to directly modulate intralimbic neural connectivity in listeners. Affective signals generally intend to influence the cognitive and neural state of listeners, and affective speech can meaningfully influence the synchronization of amygdala and hippocampal functions. Stimulating the amygdalo-hippocampal circuit further stimulates neural affective voice processing in the broader brain network, with important contributions of auditory cortical resources. Successful affective communication finally leads to a speaker-listener coupling by connecting voice production with brain decoding parameters.

## DATA AVAILABILITY

According to the Swiss Human Research Act, we cannot openly share participants’ and actors’ experimental acoustic and brain data without the explicit consensual agreement of study participants, which was not the case for the full study sample. However, data can be shared upon reasonable requests to the corresponding authors in consultation with the local ethics committee of the Cantone Zurich. Regarding the acoustic data, examples of the live affective speech can be found here: caneuro.github.io/blog/2024/study-live-voice-a2h. Regarding data from the group level analysis, the group AMY-aHC-connectivity intensity maps as well as the group T-maps of the fMRI data are available on NeuroVault (neurovault.org/collections/ZYMJLUEL) and the MNI coordinates of all significant peak voxels are listed in Table S1. For the GCA, the resulting group-level connectivity parameters between all ROIs are listed in Table S2.

## CODE AVAILABILITY

No custom-made code was used for the analyses in this study. We used standard codes implemented in the Statistical Parametric Mapping software (SPM12; RRID:SCR_007037; fil.ion.ucl.ac.uk/spm) for analysis of the fMRI data. For the Granger causality analysis, we used the MVGC Multivariate Granger Causality Matlab toolbox (mvgc_v1.0; RRID:SCR_015755; users.sussex.ac.uk/∼lionelb/MVGC) with default settings. The AMY-aHC voxel correlation, perceptual rating, acoustic feature, and correlation analyses were computed with standard Matlab functions (2018b; RRID:SCR_001622; mathworks.com/de/products/matlab) for all statistical steps that are extensively detailed in the Methods.

## ACKNOWLEDGEMENTS

This study was supported by the Swiss National Science Foundation (SNSF 100014_182135/1 to SF). We thank Numan Korkut and Caitlyn Trevor for their help in recruiting participants and collecting data. We also thank Philipp Stämpfli for technical assistance with the neurofeedback setup.

## AUTHOR CONTRIBUTIONS

F.S. contributed to designing the experiment, data acquisition, data analysis, and writing and editing the manuscript; S.F. contributed to designing the experiment, data analysis, and writing and editing the manuscript.

